# Female reproductive tract has low concentration of SARS-CoV2 receptors

**DOI:** 10.1101/2020.06.20.163097

**Authors:** Jyoti Goad, Joshua Rudolph, Aleksandar Rajkovic

## Abstract

There has been significant concern regarding fertility and reproductive outcomes during the SARS-CoV2 pandemic. Recent data suggests a high concentration of SARS-Cov2 receptors, *ACE2* or *TMPRSS2*, in nasal epithelium and cornea, which explains person-to-person transmission. We investigated the prevalence of SARS-CoV2 receptors among reproductive tissues by exploring the single-cell sequencing datasets from uterus, myometrium, ovary, fallopian tube, and breast epithelium. We did not detect significant expression of either *ACE2* or *TMPRSS2* in the normal human myometrium, uterus, ovaries, fallopian tube, or breast. Furthermore, none of the cell types in the female reproductive organs we investigated, showed the co-expression of *ACE2* with proteases, *TMPRSS2*, Cathepsin B (*CTSB*), and Cathepsin L (*CTSL*) known to facilitate the entry of SARS2-CoV2 into the host cell. These results suggest that myometrium, uterus, ovaries, fallopian tube, and breast are unlikely to be susceptible to infection by SARS-CoV2. Our findings suggest that COVID-19 is unlikely to contribute to pregnancy-related adverse outcomes such as preterm birth, transmission of COVID-19 through breast milk, oogenesis and female fertility.

## INTRODUCTION

The coronavirus disease 2019, also commonly known as COVID-19 is caused by severe acute respiratory syndrome coronavirus 2 (SARS-CoV2). SARS-CoV2 is a single-stranded positive sense RNA virus first detected in Wuhan, China in late 2019 (1, 2). Since then, it has spread worldwide, becoming a global pandemic, infecting nearly 3.58 million people worldwide and resulting in 247,503 deaths (3). The severity of SARS-CoV2 varies as infected individuals can be either asymptomatic or present mild to severe symptoms. Some of the most common symptoms presented among the individuals infected with SARS-CoV2 include fever, cough, pneumonia, occasional diarrhea, muscle pain, and new loss of sense of smell or taste (4).

SARS-CoV2 binds to angiotensin-converting enzyme 2 (*ACE2*) receptor on the host cells through spike (S) protein on the surface of the virus (2, 5). In addition to *ACE2*, entry of the virus into the host cell is also mediated by proteases *TMPRSS2* (5). In the absence of *TMPRSS2*, SARS-CoV2 is known to use cathepsins, *CTSB* and *CTSL* as an alternate to enter the host cells (5). These proteases are required for the priming of the S protein after it binds to the *ACE2* receptor for its entry into the host cell (5, 6).

Recent analyses of existing single-cell sequencing datasets showed that the SARS-CoV2 receptor, *ACE2* is expressed in various cell types of organs of the respiratory tract, with relatively high expression in goblet cells and ciliated cells of nasal epithelium and club cells in the lung (7). In addition to respiratory tract, additional single cell sequencing analyses of cornea, ileum, colon, heart, and gallbladder besides the respiratory tract identified cells that are susceptible to SARS-CoV2 infection (7). These findings may explain cardiovascular inflammation, conjunctivitis and diarrhea as well as other symptoms among individuals infected with SARS-CoV2.

One of the major clinical concerns is the effect of SARS-CoV2 on pregnancy and fertility. Reports with data from a limited number of pregnant women suggest that SARS-CoV2 is responsible for miscarriages, preterm birth, stillbirth, and fetal growth restrictions due to placental abnormalities (8, 9). However, the susceptibility of female reproductive organs to SARS-CoV2 is poorly understood.

We investigated the cell-specific presence of *ACE2/TMPRSS2* receptor expression in the female reproductive organs as a surrogate for their susceptibility to SARS-CoV2. We examined the myometrium, uterus, ovary, fallopian tube, and breast single-cell RNA sequencing datasets for cell specific expression of the SARS-CoV2 receptor, *ACE2*. Our study gave us critical insights into the expression of SARS-CoV2 receptor and proteases *TMPRSS2, CTSB/L* in the female reproductive tract. Our findings suggest that ovary, fallopian tube, uterus, myometrium, and breast are unlikely to be direct targets for SARS-CoV2 entry.

## METHODS

### Datasets and analyses

The published datasets can be found at: fallopian tube (GSE139079), breast (NCBI GSE113197), ovary (https://www.ebi.ac.uk/arrayexpress/experiments/E-MTAB-8381/), and uterus (GSE134355). We retained the cell clustering the same as described in the respective manuscripts except for the uterus. For the uterus dataset, we filtered out cells expressing more than 20% mitochondrial genes and used the standard Seurat pipeline to obtain the cell clusters.

### Myometrium tissue collection and preparation of the single-cell suspension

Normal myometrium was collected from the patients undergoing hysterectomy with informed consent. Tissues were collected under the UCSF Biospecimen Resources (BIOS) program approved by the institutional review board, ethics approval, 17-22669.

Fresh tissue samples were collected, stored in ice-cold HBSS, and transported to the lab. The myometrium was cut into 3-4 mm pieces. These pieces were then added to the 3-4 ml of digestion media containing 0.1 mg/ml liberase (Roche, 501003280), 100 U/ml DNase I (Sigma, D4527), and 25 U/ml dispase (Sigma, D4818) in DMEM (Life Technologies, 12634010) per gm of the tissue, and mechanically dissociated using the gentle MACS dissociator (Miltenyi Biotech) for 30 mins at 37°C to prepare the single-cell suspension. The cell suspension was then pipetted up and down with 25 ml,10 ml, and 5 ml pipette for 1 minute each and then filtered through 70 μm filter (Corning, 431751). Debris was then removed from the cell suspension using the debris removal solution (Miltenyi Biotech, 130-109-398) as per the manufacturer’s instructions. The cells were then incubated with RBC lysis buffer (Thermofisher Scientific, 00-4333-57) for 5 mins on ice to remove the red blood cells. The cells were then resuspended in the 0.4% ultrapure BSA (Thermofisher Scientific, AM2616) in PBS and passed through the 70-μm cell strainer (Bel-Art, H13680-0070) to obtain the single-cell suspension.

### Single-cell RNA sequencing library preparation

Single cells were processed through 10X Chromium system (10X Genomics, USA) using the single-cell 5’ library and the gel bead kit (10X Genomics, PN-1000006), and the Chromium single cell chip kit (10X Genomics, PN-1000151) as per the manufacturer’s instructions. The cells were partitioned into barcoded gel bead-in-emulsions reverse transcription was performed on individual droplets. cDNA libraries were then sequenced using an Illumina Hi-Seq 2500/NOVA (Illumina).

### Single-cell RNA-seq data preprocessing and analysis for myometrium dataset

FASTQ files were analyzed using Cellranger (version 3.1; 10x Genomics). Raw count cell by transcript matrices were imported into R and Seurat (version 3.0) and used for further analysis. For quality control, cells having fewer than 200 reads, greater than 2500 reads, or more than seven percent mitochondrial gene expression were removed. The “sctransform” function was utilized to integrate the Seurat object from the eight tissue samples. Any cells with more than one percent expression of hemoglobin genes *HBA2, HBA1*, and *HBB* were removed. The raw transcripts were normalized in each cell to transcripts per 10,000 UMI to remove the batch effects and log2 transformed. The uniform manifold approximation and projection (UMAP) was used for dimensionality reduction and clustering the cells.

## RESULTS

### Expression of ACE2 or TMPRSS2 in the ovary

Normal ovarian function is essential for proper oogenesis and fertility. We investigated the presence of the *ACE2, TMPRSS2, CTSB*, and *CTSL* in the human ovary cell types derived from single-cell sequencing (10). Data processing and cluster annotation were performed as previously described (10) (Fig. 1A, B). We found that *ACE2* was expressed at a very low level in less than 5% of stroma and perivascular cells of the ovarian cortex. We did not observe the expression of *TMPRSS2* in any of the eight distinct cell types in the ovary (Fig. 1C, D). *CTSB* and *CTSL* were found to be expressed in all eight ovarian cell types (Fig. 1C, D). However, we did not observe any cells in the ovary co-expressing *ACE2/ CTSB or ACE2/CTSL* (Fig. 1D). Since *ACE2* requires the co-expression of protease *TMPRSS2* or *CTSB*/L to facilitate its entry into the host cell by priming the S protein on its surface, our data suggest that SARS-CoV2 is unlikely to infect the ovarian cells and unlikely to affect oogenesis.

**Fig1.**
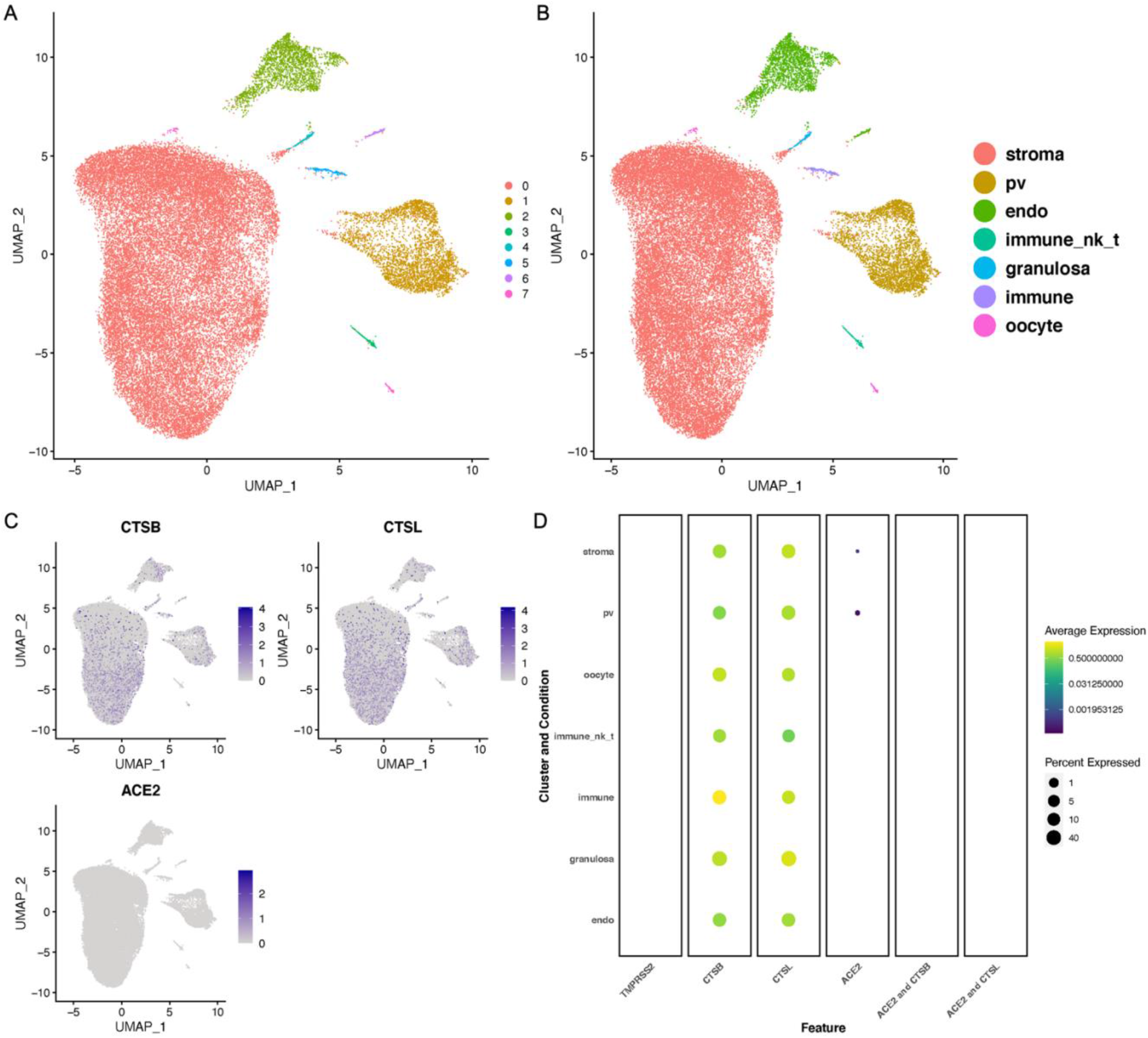
Expression of *ACE2*, and cathepsins B and L in the human ovary: A) UMAP showing the number of different clusters in ovary. B) UMAP projection with the cell annotations in the human ovarian cortex. Pv, perivascular cells; endo, endothelial cells; immune NK-T, natural killer cells and T-cells. The raw data was normalized, log transformed and analyzed same as the described in the published paper (10). C) Feature plots showing the expression of the *ACE2, TMPRSS2, CTSB*, and *CTSL* in the ovary UMAP, grey: No RNA expression purple: RNA positive. D) Dot plots showing the expression of the genes in each cell type along with the co-expression of *ACE2/CTSB* and *ACE2/CTSL* (with Benjamini Hochberg adjusted p value). The dot size represents the proportion of the cells within the respective cell type expressing the gene and the color indicates the average gene expression.

### Expression of *ACE2* and proteases *TMPRSS2, CTSB*/L in the fallopian tube

The fallopian tube is responsible for transporting the oocyte or fertilized egg to the uterus for implantation. Inflammation and blockage of the fallopian tube can lead to infertility in women. We therefore analyzed SARS-CoV2 receptors expression in different cell types of the fallopian tube. We analyzed the previously published single-cell dataset from the normal fallopian tube (11). We performed the cluster annotation and filtering of the low-quality cells exactly as previously described (11). Cell clusters and cluster annotations are shown in Fig. 2A, B. Our analysis of the fallopian tube dataset revealed *ACE2* expression in less than 5% ciliated cells, secretory cells, and leukocytes. In contrast, proteases *TMPRSS2* and *CTSL/B* showed varying expression levels in all cell types of oviduct. We did not detect expression of *CTSL* in any cell type of fallopian tube (Fig. 2C, D). We did not observe any fallopian tube cells co-expressing *ACE2* with either of *TMPRSS2* or *CTSB*. We know that the ciliated cells in the fallopian tube are essential for the movement of oocyte and sperm through the lumen, while secretory cells secrete nutrient-rich fluid for the egg and sperm to find each other (12–14). Together, this data suggests that SARS-CoV2 infection is unlikely to affect early fertilization events.

**Fig2.**
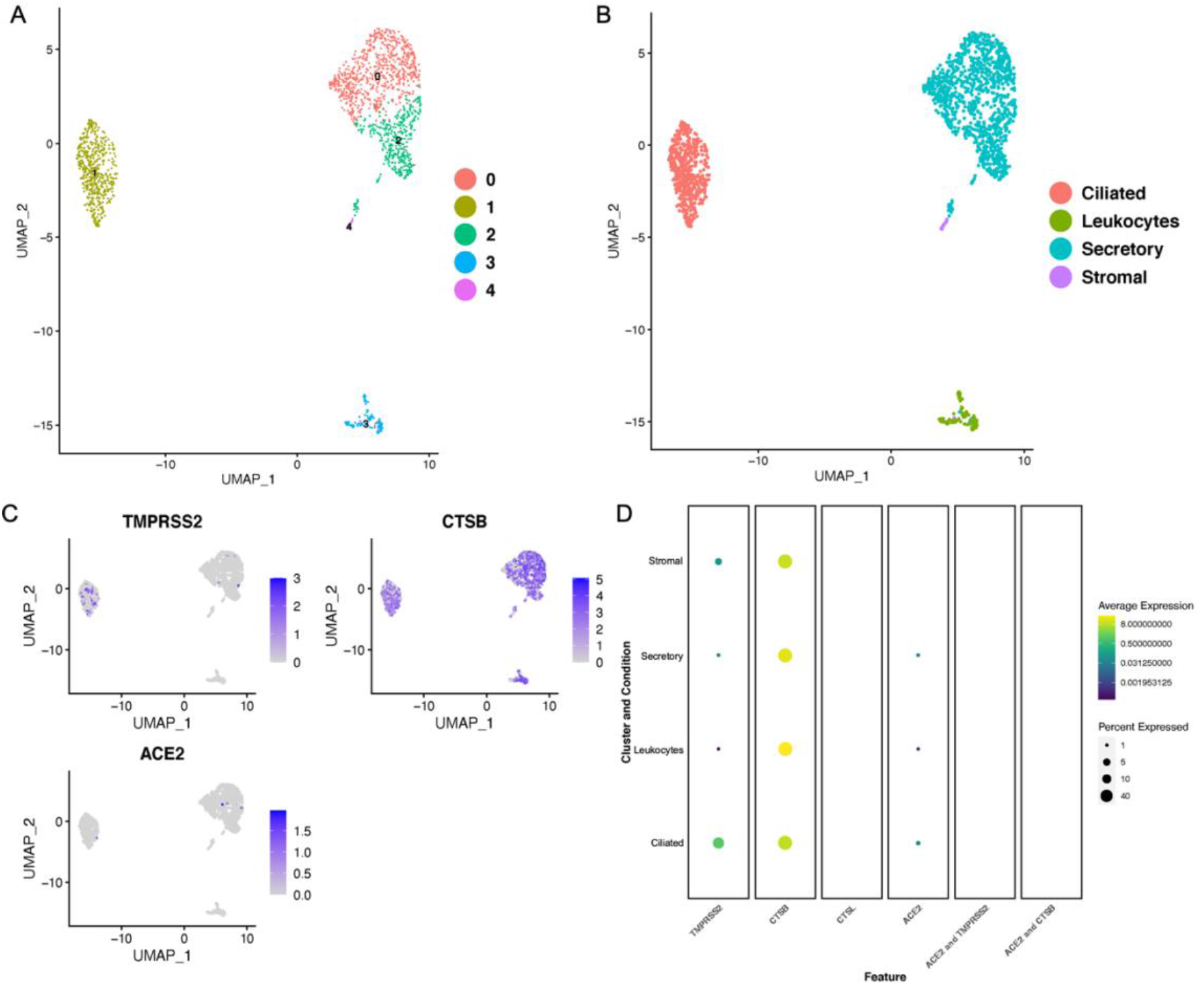
Expression of *ACE2* and proteases *TMPRSS2, CTSB*/L in the fallopian tube. A) UMAP projections of the cell clusters in normal fallopian tube B) UMAP showing the cell annotation of the normal fallopian tube. C) Feature plots showing the expression of SARS-CoV2 receptor, *ACE2*, and proteases *TMPRSS2, CTSB* and *CTSL* in the normal fallopian tube grey: No RNA expression purple: RNA positive D) Dot plots showing the expression of the genes in each cell type along with the co-expression of the *ACE2/TMPRSS2* and *ACE2/ CTSB* in the normal fallopian tube (with Benjamini–Hochberg-adjusted p values). The dot size represents the proportion of the cells within the respective cell type expressing the gene and the color indicates the average gene expression.

### Myometrium expression of SARS-CoV2 receptors

We performed single-cell transcriptome analysis of the normal myometrium collected from the women undergoing hysterectomy. Seurat analysis of 11,235 high-quality cells from the myometrium revealed the presence of 13 distinct cell populations (Fig. 3A). The subpopulations included known cell types: smooth muscle cells, fibroblasts, natural killer (NK) cells, T cells, myeloid cells, and endothelial cells (Fig. 3B). These 13 subpopulations were annotated by performing the differential gene expression analysis supported by the known markers such as *ACTA2, CNN1* for smooth muscle, *VWF* and *PECAM* for endothelial cells, *DCN* and *LUM* for fibroblasts, *CD3D* for T cells, *GNLY* and *NKG7* for NK cells, *PROX1* for lymphatic endothelial cells and *CD14*, and *S100A8* for myeloid cells (S.Fig. 1). We observed low expression of *ACE2* in approximately 1% of the fibroblast cells in the myometrium. We did not observe expression of *TMPRSS2* in any of the cell types in normal myometrium (Fig. 3C,D). We also investigated the presence of cathepsins in these cell types. It has been previously reported that *CTSB* and *CTSL* are potentially involved in facilitating the entry of the SARS-CoV2 into the cell in the absence of *TMPRSS2* (5). We discovered that myeloid cells, endothelial cells, lymphatic endothelial cells, T-cells, NK cells, and fibroblast cell populations in the myometrium express *CTSB* and *CTSL* (Fig. 3C, D). Interestingly, we did not find co-expression of either *CTSB* or *CTSL* with *ACE2* (Fig. 3D). These findings indicate that SARS-CoV2 is unlikely to infect the smooth muscle cells in the myometrium, suggesting that COVID-19 infection is unlikely to cause myometrial inflammation and potentially preterm birth.

**Fig3.**
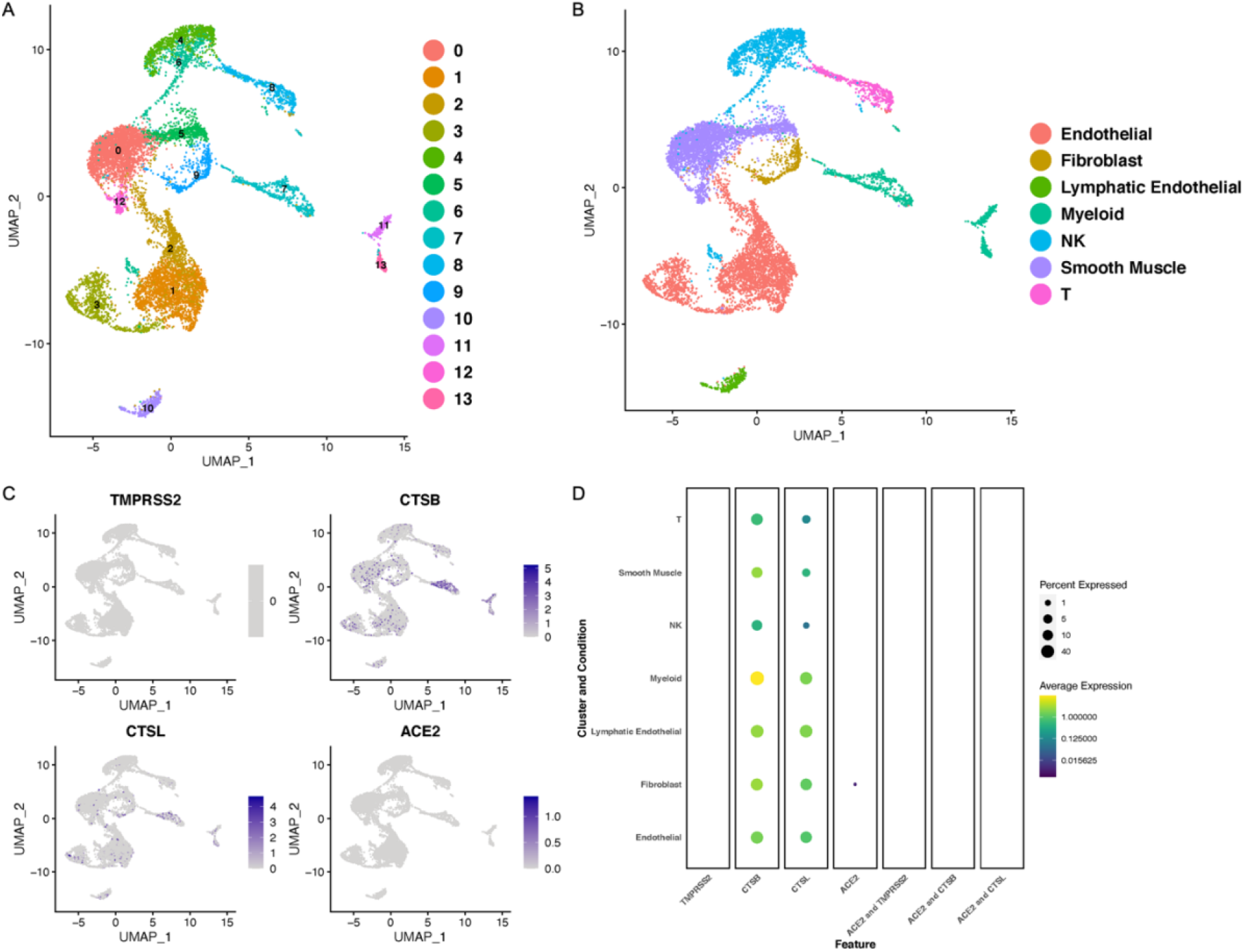
ACE2 and TMPRSS2 expression in the human normal myometrium: A) UMAP projection of the number of different cell clusters in the normal human myometrium. B) UMAP projection with the cell annotations in the normal human myometrium. C) Feature plots showing the expression of the *ACE2, TMPRSS2, CTSB*, and *CTSL* in the myometrium UMAP, grey: No RNA expression purple: RNA positive D) Dot plots showing the expression of the genes in each cell type (with Benjamini–Hochberg-adjusted p values). The dot size represents the proportion of the cells within the respective cell type expressing the gene and the color indicates the average gene expression.

### Cell-specific expression of SARS-CoV2 receptors in uterus

Once the egg is fertilized, the uterus plays a critical role in the implantation and maintenance of pregnancy. Our data in the myometrium did not indicate co-expression for genes necessary for SARS*-*CoV2 infection. To determine if SARS-CoV2 might affect the uterine function, we wanted to investigate the expression of SARS-CoV2 receptor in the whole uterus. Therefore, we investigated the co-expression of *ACE2* with *TMPRSS2* and *CTSB/L* from single-cell sequencing of the whole uterus (15). Analysis of uterine dataset revealed presence of 10 clusters including smooth muscle cells, stromal cells, luminal epithelium, endothelial cells, fibroblasts, and macrophages (Fig. 4A, B). Expression analysis of *TMRSS2* revealed the absence of *TMPRSS2* RNA in all cell types of the uterus. We found very low expression of *ACE2* in approximately 5% of the stromal cells and 1% endothelial cells (Fig. 4C, D). We also assessed the expression of *CTSB*/*L* in the uterus dataset and found that both *CTSB/L* are expressed in all cell types of the uterus (Fig. 4C, D). However, none of these cell types co-expressed *ACE2* with either *CTSL* or *CTSB* (Fig. 4D). These findings suggest that it is unlikely that uterus is susceptible to SARS-CoV2 infection in humans.

**Fig4.**
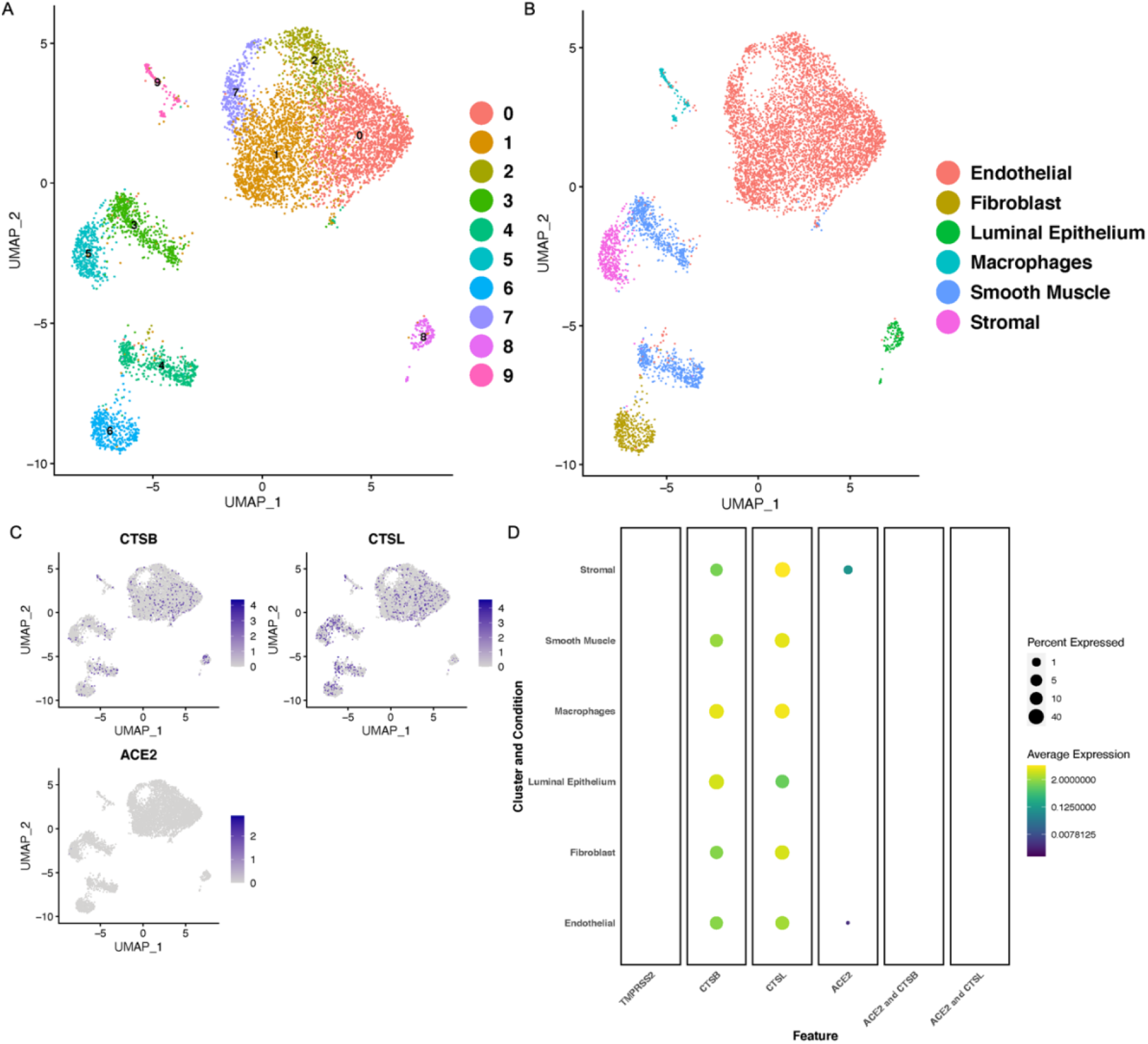
Cell-specific expression of SARS-CoV2 receptors in human uterus. A) UMAP projections of the cell clusters in normal uterus B) UMAP showing the cell annotation of the human uterus tube. C) Feature plots showing the expression of SARS-CoV2 receptor, *ACE2*, and proteases *TMPRSS2, CTSB* and *CTSL* in the uterus grey: No RNA expression purple: RNA positive D) Dot plots showing the expression of the genes in each cell type along with the co-expression of the *ACE2/TMPRSS2* and *ACE2/ CTSB* in the uterus (with Benjamini–Hochberg-adjusted p values). The dot size represents the proportion of the cells within the respective cell type expressing the gene and the color indicates the average gene expression.

### Cell-specific expression of ACE2 and TMPRSS2 in breast epithelium

We also wanted to investigate if the SARS-CoV2 can infect the mammary gland epithelium cells and potentially be transmitted to the neonates through the breast milk. We investigated the presence of *ACE2, TMPRSS2*, and *CTSBL/B* within the single-cell sequencing dataset from the primary human breast epithelial cells (16). These samples were collected from patients undergoing reduction mammoplasties. UMAP) and cell cluster annotations are shown in Fig 5A and B. We found that *ACE2* was expressed in approximately 5% of luminal epithelium and myofibroblasts in breast epithelium (Fig. 5C, D). *TMPRSS2* was expressed at very low levels in the luminal epithelium, basal and myofibroblast cells (Fig. 5C, D). Both *CTSB* and *CTSL* were expressed all cell types of the breast epithelium (Fig. 5C, D). However, we did not find any cells in the breast epithelium co-expressing *ACE2* and either of the proteases (Fig. 5D). As the co-expression of *ACE2/TMPRSS2* or *ACE2/ CTSB/L* is important for the entry of the virus into the cell, these findings indicate that there is no risk of vertical transmission of SARS-CoV2 in neonates through breastfeeding by infected mother as breast is unlikely to be infected by SARS-CoV2.

**Fig5.**
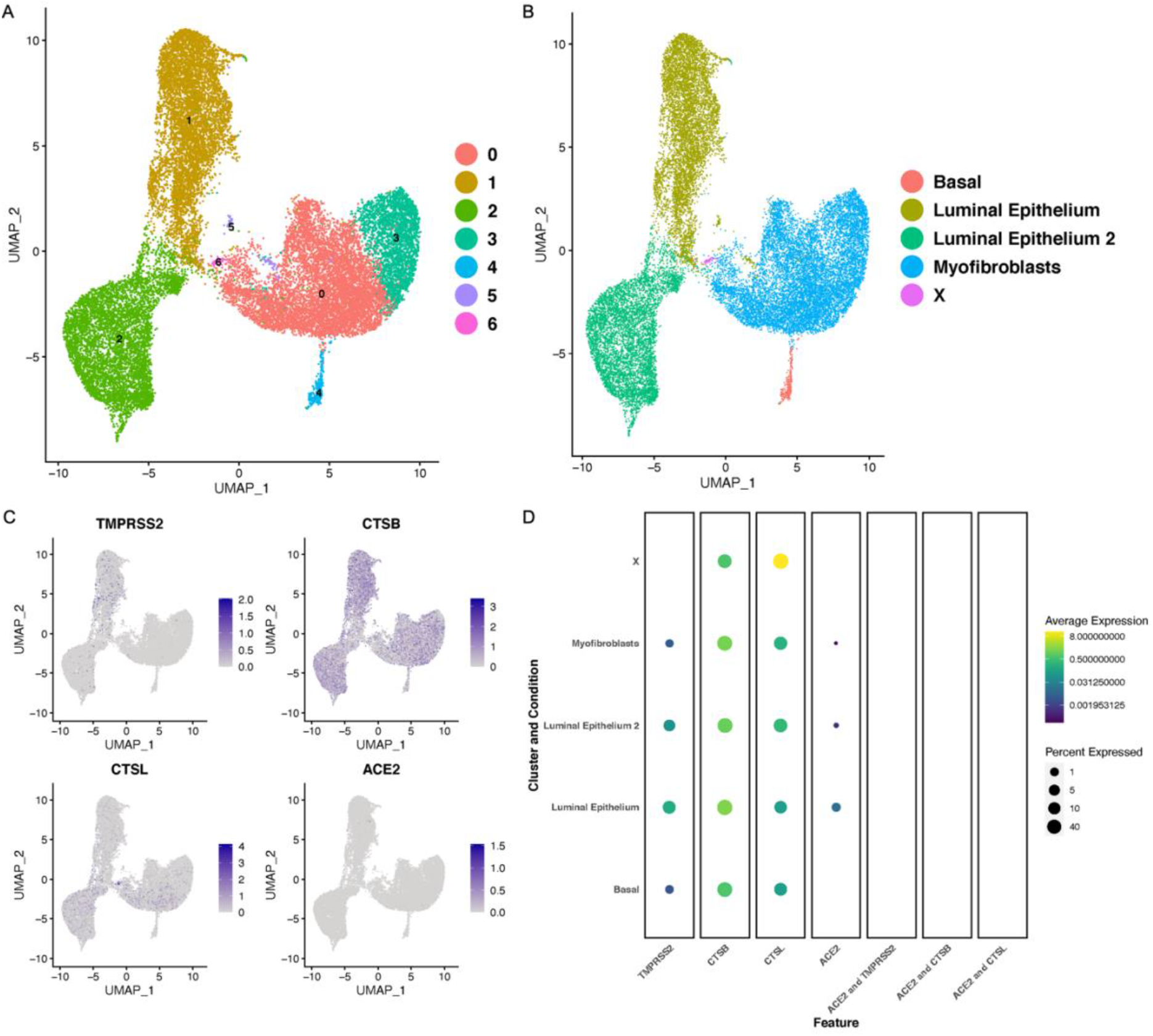
Expression of COVID receptors in the human breast epithelium: A) UMAP showing the number of different clusters in breast epithelium. B) UMAP projections with the cell annotations in the human breast epithelium. C) Feature plots showing the expression of ACE2, *TMPRSS2, CTSB* and *CTSL* in the breast epithelium. D) Dot plots showing the expression of *ACE2, TMPRSS2, CTSB*, and *CTSL* along with co-expression of *ACE2/TMPRSS2, ACE2/CTSB* and *ACE2/CTSL* in different cell clusters in the breast epithelium. The dot size is indicative of the expression of the genes in the cell type with Benjamini Hochberg adjusted p value). The dot size represents the proportion of the cells within the respective cell type expressing the gene and the color indicates the average gene expression.

## DISCUSSION

With the SARS-CoV2 infection affecting multiple organs, there are increasing concerns about the effect of SARS-CoV2 on pregnancy and fertility (17). There is very limited and conflicting data on how COVID-19 affects pregnancy and transmission of SARS-CoV2 from mother to the neonate (9). In this study, we analyzed single-cell sequencing datasets from uterus, ovary, fallopian tube, and breast to better understand the susceptibility of different cell types in the female reproductive tract to infection by SARS-CoV2.

Studies with limited patient numbers have shown that the women infected with SARS-CoV2 have a higher incidence of premature delivery, miscarriage, and intrauterine growth restriction (9). Total reports from 32 patients have suggested that 47% of women affected by COVID-19 had preterm deliveries (9). However, the aforementioned studies are small and there is no convincing evidence on whether the preterm births were directly due to SARS-CoV2 infection of the reproductive tract, secondary effects of systemic inflammation, or mechanisms unrelated to SARS-CoV2. Our data in this study revealed very low expression of *ACE2* in uterine stromal cells and endothelial cells. We did not detect expression of *TMPRSS2* in any of the uterine cell. However, *CTSB* and *CTSL* were expressed in the fibroblasts, stroma, smooth muscle cells and macrophages of the uterus. SARS-CoV2 uses *CTSB/L* proteases that act as an alternative pathway to enter the host cell in the absence of *TMPRSS2* (5). Since we did not find co-expression of *ACE2* with any of the proteases implicated in the entry of the SARS-CoV2, it seems unlikely that uterus will be affected by COVID-19. The myometrium, which regulates uterine contractions and is critical in the onset of labor, did not contain cells that co-expressed *ACE2/TMPRSS2* receptors. Together, these results indicate that COVID-19 infection is unlikely to infect myometrial cells directly. SARS-CoV2 is therefore unlikely to directly contribute to abnormal uterine function which may result in implantation failure, preterm birth, and early placentation.

We found that while *ACE2* was expressed in approximately 1% of stromal cells and perivascular cells, *TMPRSS2* was not expressed in any of the eight ovarian cell types. Furthermore, fallopian tube data showed very few ciliated cells, secretory cells, and leukocytes that expressed *ACE2*. We did not find any cells co-expressing *ACE2* and *TMPRSS2* or *CTSB/L* in either ovary or fallopian tube. Together these results suggest that SARS-CoV2 is unlikely to affect female fertility. A recent study analyzing the previously published testes single-cell sequencing dataset identified *ACE2* in spermatogonia stem cells, Leydig cells, and mast cells. However, these cells lacked co-expression of the *TMPRSS2* receptor(7). These findings together with our study suggest that the SARS-CoV2 infection is unlikely to damage fertility.

Recent studies using the already published placental single-cell sequencing datasets have shown that syncytiotrophoblast cells, villous cytotrophoblast cells, decidual perivascular cells, decidual stromal cells in placenta express *ACE2* in 6-14 weeks of gestation. However, these investigators did not observe co-expression of *ACE2* and *TMPRSS2* in any of the placental cells at this stage (7). Interestingly, another independent study found expression of *TMPRSS2* only in villous cytotrophoblast and epithelial glandular cells and syncytiotrophoblast cells, using the same dataset (Li et al. 2020). However, they also found that very few cells co-expressed both *ACE2* and TMPRSS*2* only in villous cytotrophoblast cells. They did not observe the co-expression of *TMPRSS2* and *ACE2* in any other cell types of the placenta (Li et al. 2020). In this study, the authors also compared the SARS-CoV2 receptors at different stages of placental growth and found the SARS-CoV2 receptor expression is dynamic with the placental growth, with significant increase in the expression of both *ACE2 and TMPRSS2* in the extravillous trophoblasts at 24 weeks of gestational age compared to_the early stages. The increase in the expression of the SARS-CoV2 receptor at later stages of placental development might explain the reports of the placental abnormalities leading to miscarriages or fetal growth restriction in women infected with COVID-19 (8, 18). However, further detailed investigations with large sample sizes are warranted to draw any substantial conclusions.

We also wanted to find out if breast cells are susceptible to the COVID-19 infection. Current obstetric protocols for infected mothers in labor, call for temporary separation of mother and baby to prevent SARS-CoV2 transmission. Our analysis of the mammary gland dataset revealed a low expression of *ACE2* in luminal epithelium and myofibroblasts cell types. However, we did not find co-expression of *ACE2* with *TMPRSS2 or CTSB/L* in any cell types in the breast epithelium. These findings suggest that the virus might not be able to penetrate the mammary gland cells. Therefore, the chances of transmission of the virus through breastfeeding are negligible.

Together, these results suggest that major reproductive organs involved in female fertility and pregnancy are not susceptible to direct SARS-CoV2 infection. These data may explain low incidence of complications among pregnant women and little evidence for higher infertility (17). Our analyses is limited by the current single cell sequencing data sets and somewhat limited number of cell population in each individual organ. Moreover, SARS-CoV2 systemic infection is known to affect vasculature (19) as well as increase the risk of thrombosis (20) and these abnormalities may be significantly more detrimental factors to fertility and pregnancy than direct infection on the reproductive organs. Prospective studies on couples that conceive are necessary to better define the true effect of SARS-CoV2 infection on fertility and adverse pregnancy outcomes.

## CODE AVAILABILITY

https://github.com/joshucsf/cell_specific_ace2_in_female_repro

## AUTHOR’S ROLES

JG and AR designed the study, JG and JR performed the data analysis; JG and AR wrote the manuscript. AR supervised the study and provided financial support. All authors reviewed, commented and approved the manuscript

## ACKNOWLEDGEMENTS

We thank all the authors who provided the online-available single cell sequencing data used in this study. This work was supported by funding from National Institute of Child Health and Human Development (5P50HD098580).

---

**S.Fig1.**
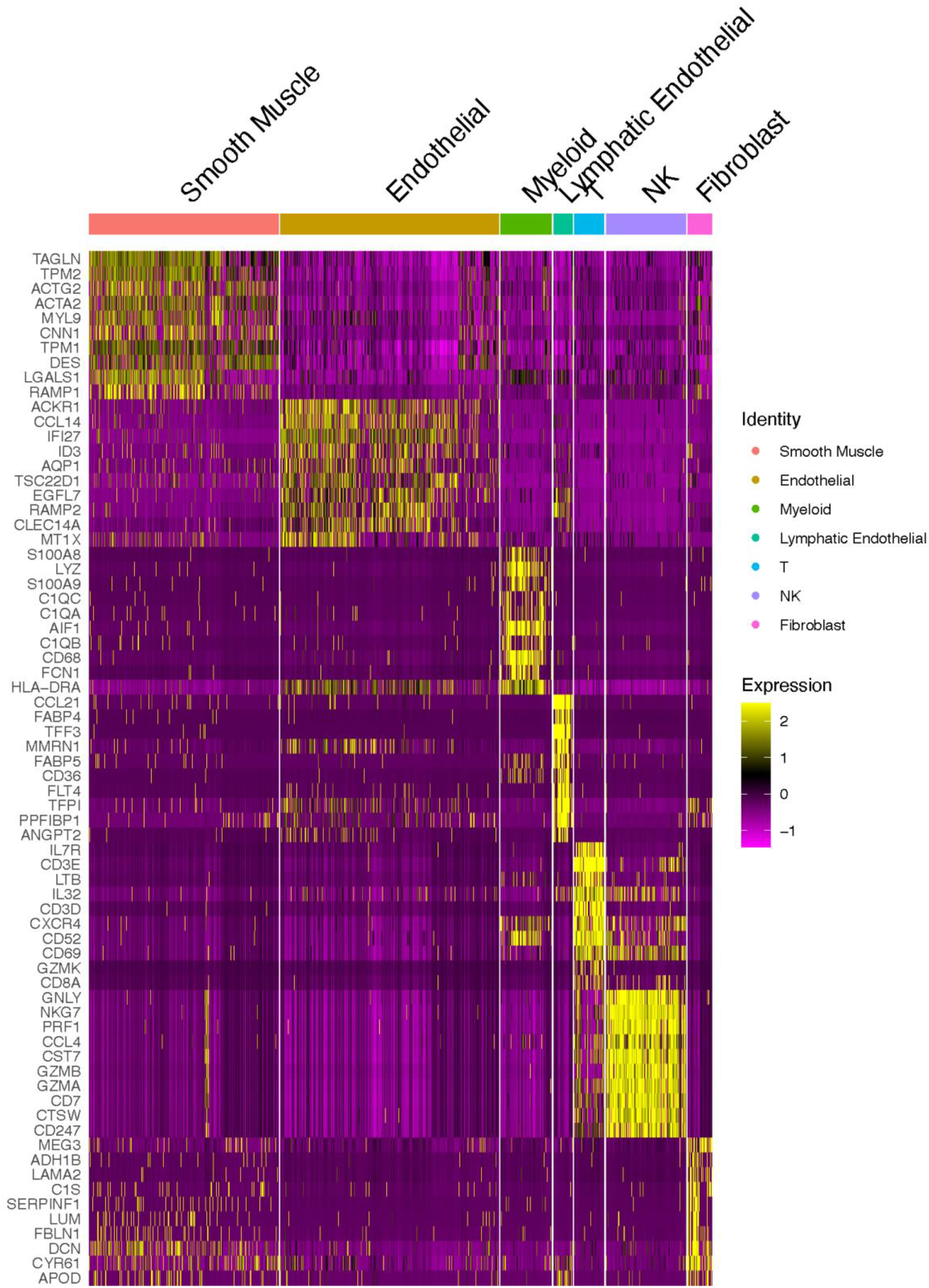
Heatmap showing the cluster annotation based on the expression of the top genes in the cell cluster. Colors represent the expression level as shown in the scale bar.

## REFERENCES

1. Fung TS, Liu DX. Human Coronavirus: Host-Pathogen Interaction. Annu Rev Microbiol 2019;73:529–57.

2. Chen N, Zhou M, Dong X, Qu J, Gong F, Han Y et al. Epidemiological and clinical characteristics of 99 cases of 2019 novel coronavirus pneumonia in Wuhan, China: a descriptive study. Lancet 2020;395:507–13.

3. . Situation report-107. In: World health organization, 2020.

4. . In: Centers for Disease Control and Prevention 2020.

5. Hoffmann M, Kleine-Weber H, Schroeder S, Kruger N, Herrler T, Erichsen S et al. SARS-CoV-2 Cell Entry Depends on ACE2 and TMPRSS2 and Is Blocked by a Clinically Proven Protease Inhibitor. Cell 2020;181:271–80 e8.

6. Iwata-Yoshikawa N, Okamura T, Shimizu Y, Hasegawa H, Takeda M, Nagata N. TMPRSS2 Contributes to Virus Spread and Immunopathology in the Airways of Murine Models after Coronavirus Infection. J Virol 2019;93.

7. Sungnak W, Huang N, Becavin C, Berg M, Queen R, Litvinukova M et al. SARS-CoV-2 entry factors are highly expressed in nasal epithelial cells together with innate immune genes. Nat Med 2020;26:681–7.

8. Shanes ED, Mithal LB, Otero S, Azad HA, Miller ES, Goldstein JA. Placental pathology in COVID-19. medRxiv 2020:2020.05.08.20093229.

9. Mullins E, Evans D, Viner RM, O’Brien P, Morris E. Coronavirus in pregnancy and delivery: rapid review. Ultrasound Obstet Gynecol 2020;55:586–92.

10. Wagner M, Yoshihara M, Douagi I, Damdimopoulos A, Panula S, Petropoulos S et al. Single-cell analysis of human ovarian cortex identifies distinct cell populations but no oogonial stem cells. Nat Commun 2020;11:1147.

11. Hu Z, Artibani M, Alsaadi A, Wietek N, Morotti M, Shi T et al. The Repertoire of Serous Ovarian Cancer Non-genetic Heterogeneity Revealed by Single-Cell Sequencing of Normal Fallopian Tube Epithelial Cells. Cancer Cell 2020;37:226–42 e7.

12. Paik DY, Janzen DM, Schafenacker AM, Velasco VS, Shung MS, Cheng D et al. Stem-like epithelial cells are concentrated in the distal end of the fallopian tube: a site for injury and serous cancer initiation. Stem Cells 2012;30:2487–97.

13. Afzelius BA, Camner P, Eliasson R, Mossberg B. Kartagener’s syndrome does exist. Lancet 1978;2:950.

14. Leese HJ, Tay JI, Reischl J, Downing SJ. Formation of Fallopian tubal fluid: role of a neglected epithelium. Reproduction 2001;121:339–46.

15. Han X, Zhou Z, Fei L, Sun H, Wang R, Chen Y et al. Construction of a human cell landscape at single-cell level. Nature 2020;581:303–9.

16. Nguyen QH, Pervolarakis N, Blake K, Ma D, Davis RT, James N et al. Profiling human breast epithelial cells using single cell RNA sequencing identifies cell diversity. Nat Commun 2018;9:2028.

17. Panahi L, Amiri M, Pouy S. Risks of Novel Coronavirus Disease (COVID-19) in Pregnancy; a Narrative Review. Arch Acad Emerg Med 2020;8:e34.

18. Baud D, Greub G, Favre G, Gengler C, Jaton K, Dubruc E et al. Second-Trimester Miscarriage in a Pregnant Woman With SARS-CoV-2 Infection. JAMA 2020.

19. Ackermann M, Verleden SE, Kuehnel M, Haverich A, Welte T, Laenger F et al. Pulmonary Vascular Endothelialitis, Thrombosis, and Angiogenesis in Covid-19. N Engl J Med 2020.

20. Connors JM, Levy JH. COVID-19 and its implications for thrombosis and anticoagulation. Blood 2020;135:2033–40.

